# NemaLife: A structured microfluidic culture device optimized for aging studies in crawling *C. elegans*

**DOI:** 10.1101/675827

**Authors:** Mizanur Rahman, Hunter Edwards, Nikolajs Birze, Rebecca Gabrilska, Kendra P. Rumbaugh, Jerzy Blawzdziewicz, Nathaniel J. Szewczyk, Monica Driscoll, Siva A. Vanapalli

## Abstract

*Caenorhabditis elegans* is a powerful animal model in aging research. Standard longevity assays on agar plates involve the tedious task of picking and transferring animals to prevent younger progeny from contaminating age-synchronized adult populations. Large-scale studies employ progeny-blocking drugs or sterile mutants to avoid progeny contamination, but such manipulations change adult physiology and alter the influence of reproduction on normal aging. Moreover, for some agar growth-based technology platforms, such as automated lifespan machines, reagents such as food or drugs cannot be readily added/removed after initiation of the study. Current microfluidic approaches are well-suited to address these limitations, but in their liquid-based environments animals swim rather than crawl, introducing swim-induced stress in the lifespan analysis. Here we report a simple microfluidic device that we call NemaLife that features: 1) an optimized micropillar arena in which animals can crawl, 2) sieve channels that separate progeny and prevent the loss of adults from the arena during culture maintenance, and 3) ports which allow rapid accessibility to feed the adult-only population and introduce reagents as needed. Culture maintenance and liquid manipulation are performed with simple hand-held syringes to facilitate integration of our technology into general laboratory protocols. Additionally, device geometry and feeding protocols were designed to emulate the body gait, locomotion, and lifespan of animals reared on agar. We validated our approach with longevity analyses of classical aging mutants (*daf-2*, *age-1*, *eat-2*, and *daf-16*) and animals subjected to RNAi knockdown of age-related genes (*age-1* and *daf-16*). We also showed that healthspan measures such as pharyngeal pumping and tap-induced stimulated reversals can be scored across the lifespan. Overall, the capacity to generate reliable lifespan and physiological data from the NemaLife chip underscores the potential of this device to accelerate healthspan and lifespan investigations in *C. elegans*.

## I. Introduction

Aging is a significant risk factor for a broad range of diseases including neurodegenerative disorders, diabetes and cancer^1–5^. With the growing aging population, the socioeconomic burden attributed with age-associated diseases is staggering and development of therapies that promote healthy aging is imperative. *C. elegans* is a powerful model organism for aging investigations with a short lifespan (3-5 weeks), remarkable genetic similarity with humans (~ 38 % orthologs^6^) and conserved signaling pathways^7^. Additionally, a fully mapped genome^8^ and incredible genetic plasticity^9,10^ makes C. *elegans* an attractive tool for aging studies. Advances in fluorescent microscopy^11^ and genomic technology (RNAi, CRISPR)^12,13^ have further expanded the number of possible ways in which C. *elegans* can be used to study healthy aging.

Lifespan analysis has become a classic method for evaluating the effects of a wide variety of genes, proteins, and pharmaceutical compounds on aging and age-associated diseases. However, traditional lifespan analysis is generally low-throughput and lacks the capability of non-invasive health metric analysis. Aging assays are generally carried out with *C. elegans* reared on agar plates containing nematode growth media (NGM). During reproduction, adults must be manually transferred to new plates to separate progeny from the original sample. To reduce the need for manual transfers, many labs utilize a strong progeny-blocking drug (2’-deoxy-5-fluorouridine, FUdR) to maintain an adult-only population^14–16^. An alternative to this approach is the use of sterile mutants^17–19^.

The simplicity of using FUdR or sterile mutants has led to new technologies for large-scale lifespan analysis in *C. elegans*. A technology known as Lifespan Machine (LSM), allows for the rapid analysis of a population of thousands of animals grown on agar supplemented with FUdR and automatically captures sequential images to score animals death and determine lifespan^20^. The LSM technology has provided insights into temporal scaling of ageing dynamics^20,21^ and helped identify chemical compounds with robust longevity effects^22^. Similarly, WorMotel technology facilitates longitudinal analysis of individuals in agar-filled microfabricated well plates^23^. Despite the large-scale capacity of such technologies, the use of FUdR in LSM and WorMotel technologies is disconcerting as FUdR has been shown to activate stress response pathways^24,25^, increase fat accumulation^26^ and alter lifespan in some genotypes^24,26,27^.

Additionally, current technologies like LSM and WorMotel lack the capability to study the effects of temporary environmental manipulations on lifespan. Such manipulations at user-defined time intervals have been central to studies on dietary restriction^28^ and cognitive aging^29^. Currently, traditional studies and high-throughput lifespan technologies do not have the capacity to quickly and reversibly manipulate environmental conditions, limiting their utility to survival analysis on animals exposed to a singular environment.

In recent years, microfluidic approaches have begun to address the limitations of agar-based lifespan assays^30–34^. Several key advantages of using PDMS-based microfluidics include (i) excellent permeability to oxygen and carbon dioxide enabling animals to experience natural atmospheric conditions^35^; (ii) size-based separation of progeny using on-chip filters^30,31,33^, eliminating the need to prevent or reduce progeny production; (ii) precise temporal control of culture environment via addition or removal of reagents^31,33^; (iii) overall reduction in the number of censored worms; and (iv) optical transparency of devices to enable white light and fluorescence imaging.

Despite the significant advantages of microfluidics-based approaches, work to date has been limited. Hulme *et al*. developed a microfluidic device to house individuals in circular chambers with the capacity to remove progeny during reproduction. Using this design, Hulme *et al* showed that swim activity declines with age, underscoring the need to further assess healthspan measures beyond survival analysis^30^. Wen *et al.* used a similar approach but integrated two-layer valves to immobilize animals for fluorescence evaluation of oxidative stress^31^. Building on this work, Xian *et al.* developed an automated system called WormFarm with integrated computer vision algorithms to score for longevity in small scale liquid cultures ^33^. WormFarm can be used for forward and reverse genetic screens and has been applied in studies on the longevity-altering effects of glucose supplementation previously conferred on agar. Recently, Bosari *et al.* connected an array of trapping channels^36,37^ to a variant of the WormFarm device to immobilize different-aged animals and image synapses on the DA9 axon^38^. Biocommunication between segregated males and hermaphrodites and the influence of male presence on development and aging has also been explored using microfluidics^34^.

From the above-mentioned studies, it is clear that innovative microfluidic technologies have proven to be valuable tools in C. *elegans* aging research. However, there are several drawbacks to the microfluidic approaches described above. Housing worms in liquid culture for a significant portion of their lifespan has been shown to induce significant gene expression changes^39,40^. Continuous swimming results in the activation of stress response pathways followed by significant changes in gene expression. Additionally, obligated swimming in liquid culture has been shown to induce fatigue and oxidative stress, resulting in adverse effects on worm health^40–42^. In addition to swim stress, removing progeny from a liquid microfluidic environment can be ineffective and may induce injuries as all the animals are pushed against sieve channels. The presence of large numbers of progeny and worm debris can also subject sieve channels to clogging, resulting in the build-up of debris that can lead to contamination by progeny and bacteria^31,33^. In contrast to swim chambers, some studies reported micropillar chambers for *C. elegans* assays^43,44^, but none have been configured and validated for lifelong or aging investigations.

Here we report a new microfluidic technology, termed NemaLife, for aging studies in *C. elegans* that addresses the limitations of agar-based studies and current microfluidic technologies (see Table 1). NemaLife is a simple and cost-effective platform that integrates an optimized micropillar arena enabling animals to adopt a crawling gait similar to that of worms on agar medium while also acting as a sieve to retain adults and prevent fluid-induced injury when removing progeny. Use of simple hand-held syringes rather than complex microfabricated valves makes NemaLife accessible for research laboratories and reduces error conferred by multi-step fabrication. Furthermore, we show that the use of a micropillar housing arena increases the efficiency of scoring various health span metrics, including pharyngeal pumping and locomotory phenotypes compared to other microfluidics-based studies^30–33^. Validation of NemaLife through the analysis of established aging mutants, RNAi studies and various culture conditions demonstrates that this new technology provides a simple platform advancing our fundamental understanding of genetic and environmental regulators of healthy aging.

**Table 1:** Comparative analysis of the available technologies for lifespan measurement in *C. elegans*.

| Platform/Technology | FUdR | Animal gait | Validation |  |  | Pharyngeal pumping | Reference |
| --- | --- | --- | --- | --- | --- | --- | --- |
|  |  |  | Feeding optimization | RNAi efficacy | lifespan influencing mutation |  |  |
| Agar-based |  |  |  |  |  |  |  |
| Standard NGM-plate | × | crawling | NA | √ | √ | √ |  |
| Lifespan machine (Stroustrup *et. al* ) | √ | crawling | NA | √ | √ | × | 20 |
| Worm Motel (Churgin *et.al.* ) | √ | crawling | NA | √ | √ | × | 23 |
| Other culture system |  |  |  |  |  |  |  |
| 96-microtiter plate (Slolis *et. al.*) | √ | swimming | √ | × | × | × | 45 |
| Worm corrals (Pittman *et. al.*) | √ | crawling | NA | × | × | × | 19 |
| Microfluidics |  |  |  |  |  |  |  |
| Microfluidic Device (Dong *et.al.* ) | × | swimming | √ | × | × | × | 34 |
| Microfluidic Device (Wen *et.al.* ) | × | swimming | × | × | × | × | 31, 32 |
| WormFarm (Xian *et.al.* ) | × | swimming | × | √ | × | × | 33 |
| Microfluidic device (Hulme *et. al.* ) | × | swimming | × | × | × | × | 30 |
| NemaLife | × | crawling | √ | √ | √ | √ |  |

## II. Results and Discussion

### A. Optimization of the NemaLife device-design and culture conditions

*C. elegans* lifespan measurement in a chip environment requires optimization of environmental conditions to limit triggering of stress resistance pathways that can influence lifespan and alter longevity. Given that the microfluidic environment is not yet a standard laboratory environment for *C. elegans* culture and lifespan assessment, our efforts were focused on device design and culture conditions that yield reproducible lifespans that match those of *C. elegans* reared on NGM plates.

#### Basic device design

We designed the NemaLife culture device (Fig. 1 a-d) based on the criteria that it includes (i) a means to introduce young adult animals and house the growing animals in the device until their death; (ii) the capacity to effectively remove progeny while retaining adults; (iii) a design that facilitates a crawling gait similar to animals moving on agar plates; and (iv) the ability to add food and/or reagents at user-defined times. These criteria were achieved by designing a worm habitat chamber that contains a micropillar lattice (Fig. 1b,c) that allows worms to crawl rather than swim, thereby eliminating swim-induced stress^40^. Sieve channels (Fig. 1d) on the sides of the habitat chamber (see red arrows in Fig. 1b), prevents young adults from escaping the arena but allows efficient passage of eggs, larval-stage animals (L1, L2) and bacterial debris. Adjacent to the sieve channels, two side-ports (see black arrows in Fig. 1b) enables progeny washing, introduction of reagents or food, and purging of air bubbles trapped within the channels or micropillar arena. Fluid manipulation is performed manually using hand-held syringes to make the device accessible to a wide range of laboratories. Worm habitat chambers are designed to have an approximate footprint of ≈ 60 mm^2^ for a population size of 10-15 animals (4-6 mm^2^ per animal), compared to an average footprint of 2828 mm^2^ in standard studies on agar for 30 − 50 animals (60-90 mm^2^ per animal). We were able to accommodate nine of these chambers on a 50×75 mm^2^ glass slide (Fig. 1a) that could produce survival data on ≈ 100 animals.

**Fig 1:**
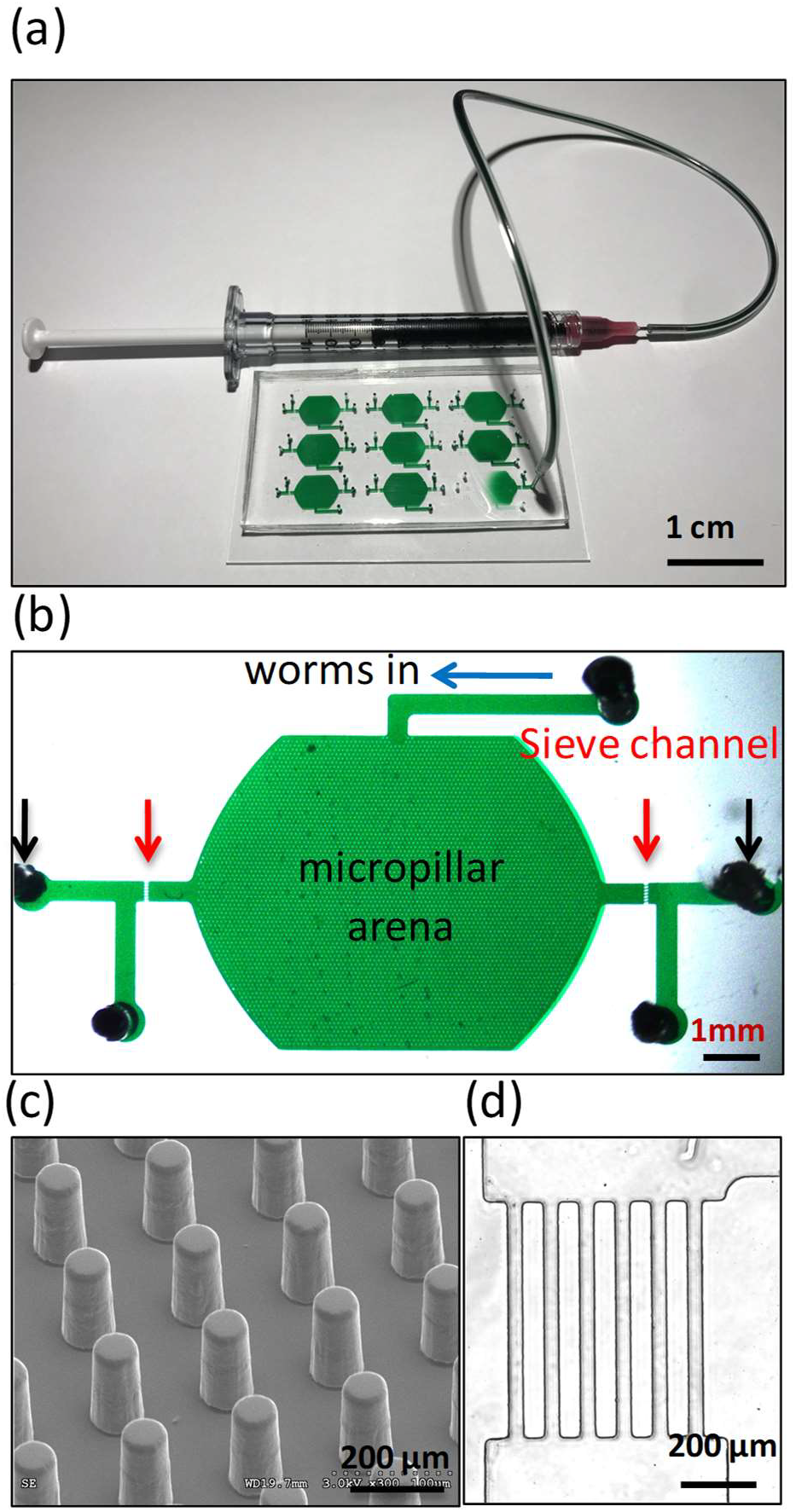
Basic design and description of the NemaLife device. (a) A 9-chamber microfluidic device for lifelong studies of crawling *C. elegans*. Fluid manipulation is performed with 1-mL syringes connected to the ports in the device. Scale bar 1 cm. (b) Design and features of the NemaLife device. Habitat arena (green) is composed of micropillars. Channel at the top (blue arrow) is the worm loading port, the two red arrows on both sides of the device identifies the sieve channel that retains worms, two black arrows indicate the reagent exchange ports and the two side ports are for air purging. Scale bar 1 mm. (c) An enlarged view of the micropillars and their lattice arrangement. Scale bar 200 µm. (d) The sieve channels consist of rectangular barriers 750 µm × 75 µm separated by a gap of 25-30 µm. Scale bar 200 µm.

The geometry of the micropillar lattice in the NemaLife chamber is crucial for successful measurement of lifespan of *C. elegans*. The lattice structure needs to accommodate significant individual variations in growing body size during reproduction and aging, maintain the natural crawling gait of *C. elegans*, and allow removal of progeny while retaining the young adults. To identify the optimal micropillar lattice that simultaneously meets these requirements, we fabricated devices with square arrangement of pillars with different pillar diameter (*a*) and gap (*s*). The nominal dimensions we tested are: Device I, *a* = 40 μm, *s* = 60 μm; Device II, *a* = 50 μm, *s* = 80 μm and Device III, *a* = 60 μm, *s* = 100 μm. The measured dimensions of the three pillar devices are reported in Table S1. During the greatest period of growth, C. *elegans* body diameter varies from ~50 − 100 µm and length varies from ~900-1500 µm. Thus, Device I provides the tightest, and Device III the leanest, confinement for the animals during the lifespan measurement. In all the devices, the pillars had a uniform height of ≈ 75 μm and a clearance from the floor of the habitat chamber of approximately ≈ 25 μm, allowing pillars to be moved aside by the animal to adjust gait and accommodate further growth. These pillar geometries and clearance facilitates progeny removal and makes it difficult for adults to escape during washing. After some initial trials, we incorporated a sieve channel design (Fig. 1d) containing rectangular blocks of 750 μm×100 μm and separated by a gap of 25-30 μm in the three devices to effectively prevent accidental removal of adults.

#### Optimization of worm culture conditions

We established culture maintenance protocols to achieve robust and reproducible lifespan data while increasing overall efficiency of aging studies. Initially, we focused on identifying the optimal washing conditions needed to remove all progeny. To do this, we cultured reproductive adults for 24 hours within the NemaLife device to allow progeny production and growth. Progeny were removed manually using gentle fluid flow, using the loading port of the device in 200 µL aliquots of S-complete.

Figure 2a shows the habitat chamber with adults and their progeny (see **SI movie 1**). After washing the retained adults are shown in Fig. 2b. Pillar environment helps worm adopt crawling gait inside the chamber (Fig. 2c) and stop being carried away with the flow. Even if the animal is already near the exit, sieve channel retains them inside the chamber (Fig. 2d). Fig. 2e shows the fraction of progeny removed (solid symbols) and the number of adults retained (open symbols) plotted against the wash volume. We find that no adult worms were lost, and all progeny were effectively removed using a total wash volume of 1 mL for three different loading conditions in the tightest lattice design (see **SI movie 2**). The washing operation took approximately 90 seconds. All trials were successful in removing progeny with this protocol, except when there was bagging, in which case more repeated washes (2-4 mL) were necessary to remove the bagged mother or the resulting advanced larval stage progeny. We note that during the washing process, animals in the chamber respond by exhibiting faster crawling momentarily, probably due to the stimulation by fluid forces during washing and feeding (see **SI movie 2 and 3**).

**Fig 2:**
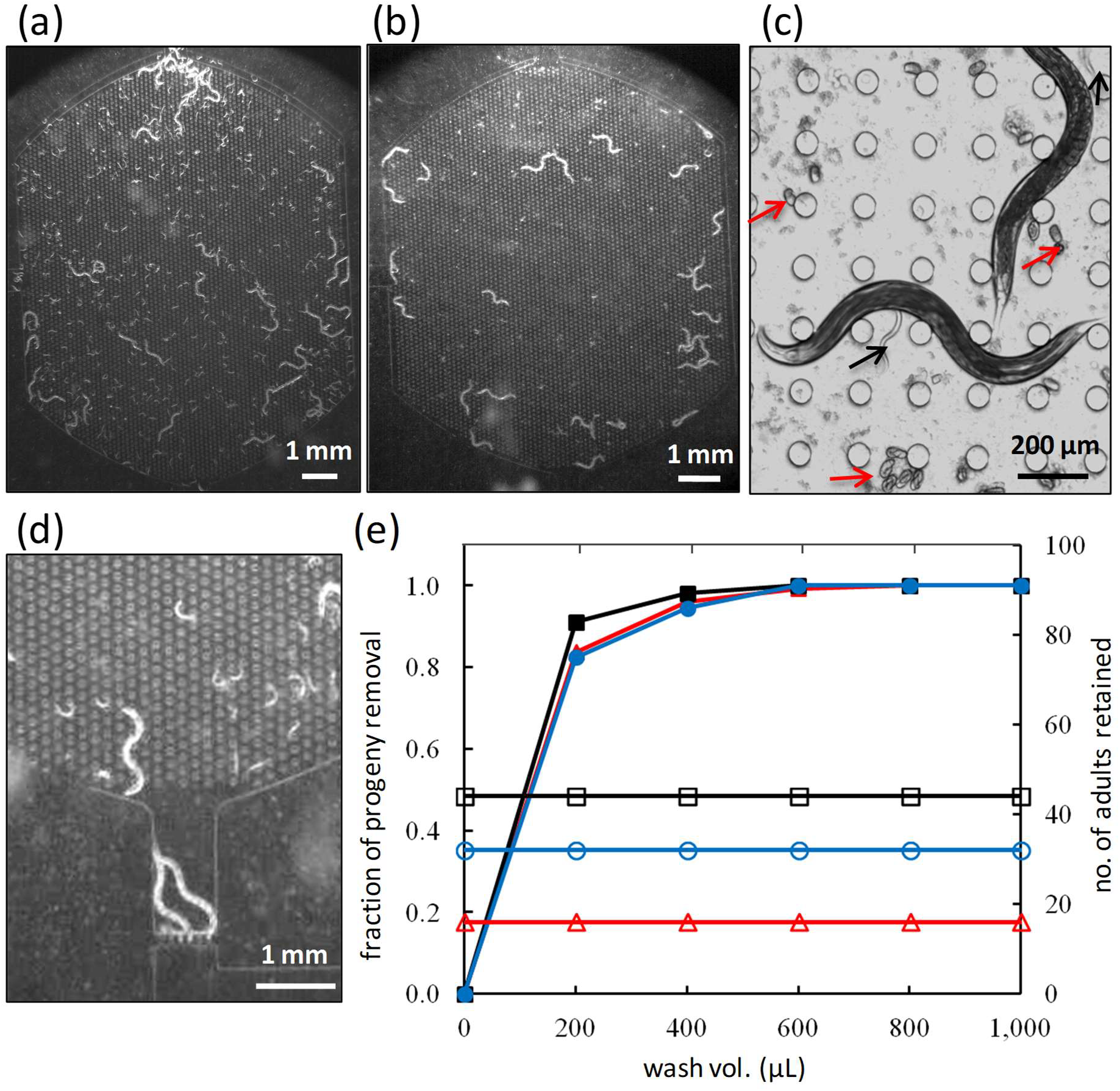
Worm culture in the NemaLife device with capacity to remove progeny. (a) A typical chamber with wild-type animals and their progeny 24 hours after loading. Scale bar 1 mm. (b) A chamber with adult-only population after removal of progeny/eggs by washing. Scale bar 1 mm. (c) Enlarged view of adult animals, progeny (black arrows) and eggs (red arrows) inside the micropillar arena. Scale bar 200 µm. (d) Animals at the exit are retained by the sieve channel, Scale bar 1 mm. (e) Effectiveness of age synchronization (progeny removal and adult retention) in NemaLife by washing the chambers with S-complete buffer. 16 (red), 32 (blue), and 44 (black) adults (day 3 after hatching) were allowed to reproduce in 3 identical units and synchronization was performed on day 4. The number of animals represents approximately 1X, 2X and 3X of the chamber capacity to check the robustness of progeny separation. Animals were incubated at 20°C. Open symbols represent adults and closed symbols represent progeny. Adult retention and progeny removal from a single unit is identified with the same color and symbol. N=3 repeat trials. The device used in efficacy trial for progeny removal has pillars of diameter 40 µm and spacing of 60 µm.

An adult *C. elegans* typically interacts with 10 − 16 pillars in the above lattice geometry design which help them to crawl normally without being washed away during fluid flow. At the same time, progeny and eggs are small enough that they can flow through the space between the pillars as well as the pillar-to-floor clearance. Pillars in NemaLife arena acts as a size based sieve. Washing experiments in Device II and III (33% and 67% more spacing between pillars than Device I) also showed that 1 mL of wash volume is sufficient for efficient clearance of progeny.

Next, we sought to establish a robust feeding protocol that generates lifespan data consistent with previous reports since specific nutrient sensing and longevity pathways can be affected by environmental conditions in addition to resource availability. *E. coli OP50* bacteria suspension is the standard diet for laboratory culture of *C. elegans* in liquid culture. Previous lifespan studies in multiwell plates^16,45^ and microfluidic devices^30,33^ used a bacterial concentration of 10^9^-10^10^ bacterial cells per mL of S-medium, a concentration equivalent to 100 mg mL^−1^ OP50^45^. In these studies, food was added to the microfluidic devices either continuously^33^ or once a day^30^. We used these previous results as a starting point to optimize the feeding protocol and evaluated the lifespan of young adult animals in the tightest geometry that were fed 100 or 200 mg mL^−1^ E. *coli* OP50 at various time intervals (Fig. 3 a,b). Worms fed intermittently (every other day) at 100 mg mL^−1^ had an overall extended lifespan and high death rate during reproduction, consistent with previous studies on intermittent fasting^46,47^. Alternatively, more frequent feeding (twice per day) did not significantly alter lifespan or age-dependent changes in body size (see SI Fig. S1). Fig. 3b shows that animals fed once a day at 100 mg/mL or 200 mg/mL had similar survival curves. However, bacterial solutions at 200 mg mL are turbid, therefore using 100 mg mL^−1^ OP50 solutions allows for better optical imaging. Thus, we settled on conducting lifespan assays in the NemaLife device by feeding 100 mg mL^−1^ of concentrated E. *coli* OP50 once every day. Additional validation studies (see Sec. II.B.) further support that this feeding regimen is optimal for *C. elegans* culture and lifespan assays in our microfluidic devices.

**Fig 3:**
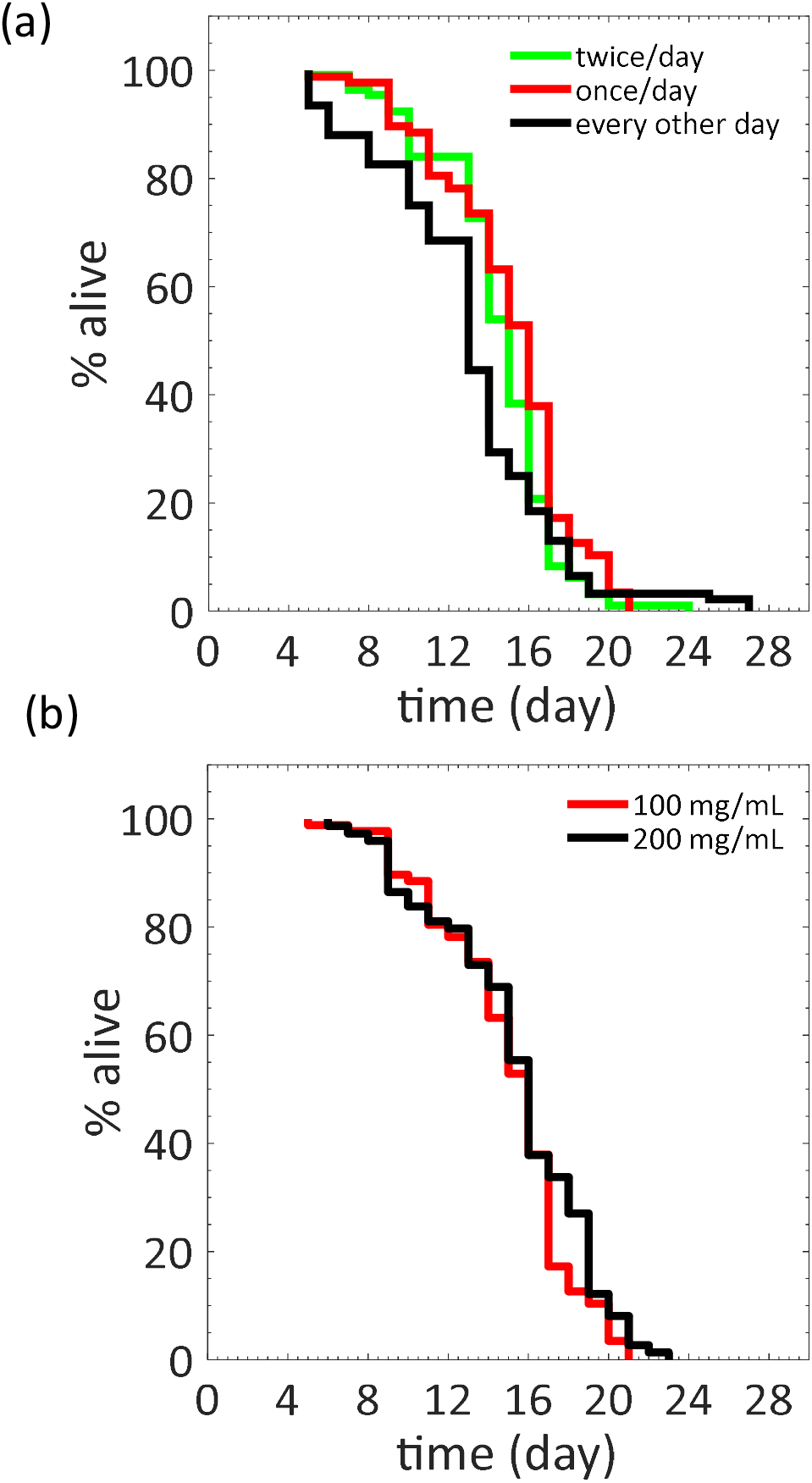
Optimization of feeding protocol for *C. elegans* lifespan assays. (a) Lifespan of wild type *C. elegans* in Device I for feeding frequency of twice per day (n=112), once per day (n=87) and every other day (n=92). N=2 repeat trials. Food concentration: 100 mg/mL of *E. coli* OP50 in S-complete. (b) Lifespan of wild type *C. elegans* for food concentration of 100 mg/mL (n=87) and 200 mg/mL (n=74) of *E.coli* OP50 in S-complete. N=1 repeat trial. Feeding frequency: once every day. Feeding animals every other day, produces a lower median (and mean) and higher maximal lifespan than feeding every day (p *=* 0.0036) or twice per day (p <0.001). P-value (once per day vs twice per day) = 0.0244.

#### Selection of the optimal micropillar arena geometry

We fabricated devices with three different micropillar arena geometries such that the gap between pillars in Devices I, II and III are *s* = 60, 80 and 100 µm, respectively. Given that the mid-length body diameter can reach as high as 95 µm in older adults (see SI Fig. S1), the animals in Device I and II are constrained, but in Device III they are not. We measured the crawling gait in terms of wavelength, amplitude and crawling speed in the three devices on day 4 and day 8. The amplitude and wavelength corresponding to the worm undulatory motion in the three devices were similar (Fig. 4a). Comparing these data with that for crawling animals on agar^48^, we find that the amplitude are similar^49^, however, the wavelength is about 30% higher on agar. We found that animal crawls in Device III with a very similar speed to that of crawling on agar surface^48^.

**Fig 4:**
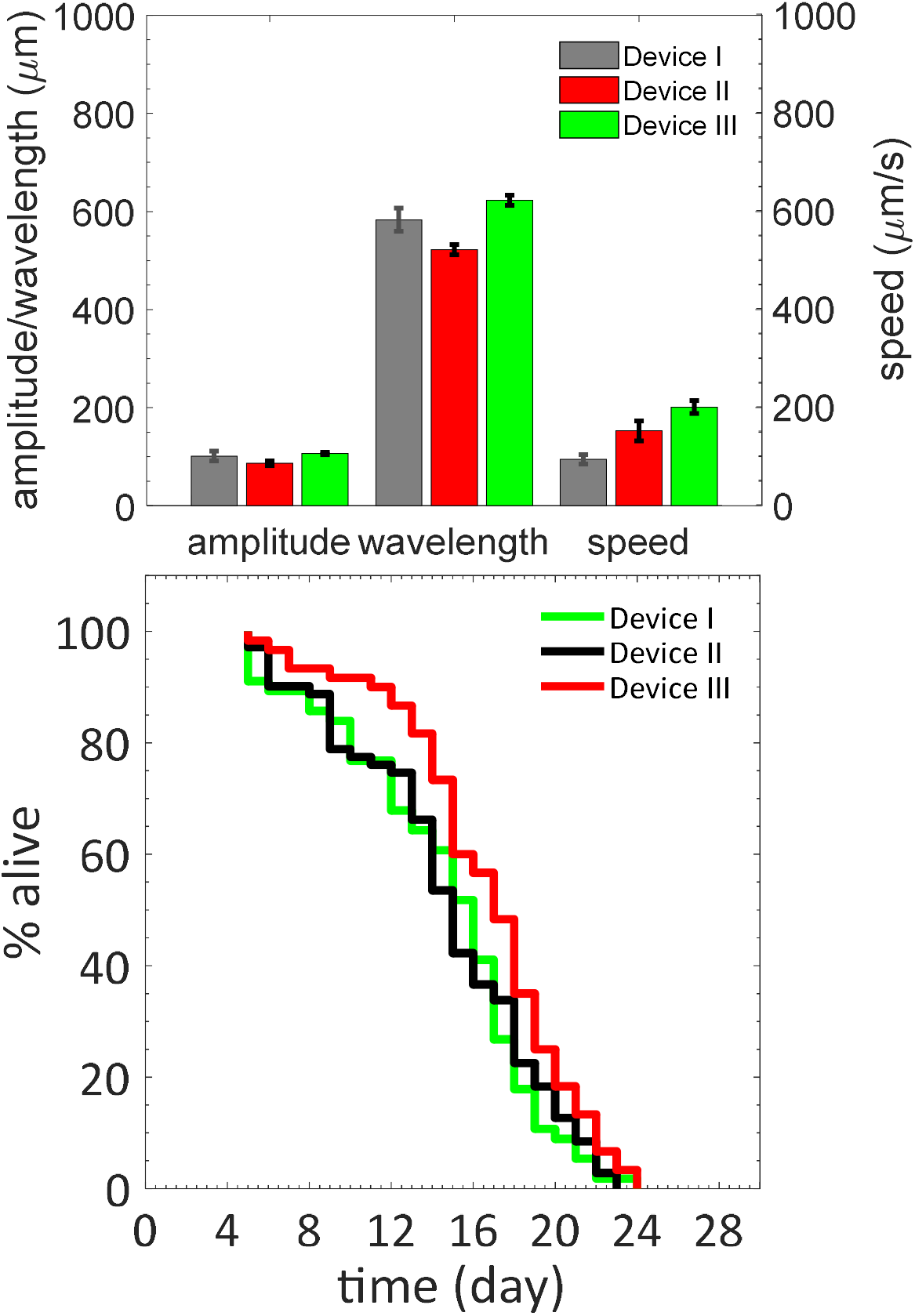
Influence of arena geometry on animal locomotion and lifespan. (a) Crawling amplitude, wavelength, and speed of day 4 animals as pillar spacing was changed within the three devices (n =10). Pillar diameter and gap for Devices I, II and III are: 40 μm, 60 μm; 50 μm, 80 μm; 60 μm, 100 μm. Older adults are most constrained in Device I and not at all in Device III. (b) Wild type *C. elegans* lifespan as a function of confinement (n=56 for Device-I, n=71 for Device-II and n=60 for Device-III). p-value for lifespan curve between Device-I and II is 0.99 and p-value for lifespan curve between Device-I and III is 0.026 (Log-rank (Mantel-Cox) test, N=2 repeat trials).

Fig. 4b shows the survival data for the animals in devices of three different geometries. We find that median and maximum lifespan is most consistent with studies on agar in device III with s = 100 µm. Devices with tighter pillar spacing resulted in a reduction of worm lifespan possibly due to restraints in natural locomotion and stress conferred through body confinement. All subsequent NemaLife aging investigations were conducted using Device III (a = 60 µm and s = 100 µm).

### B. Validation of the NemaLife device

In the previous section, we optimized the progeny removal conditions, feeding regimen and micropillar geometry to achieve efficient culture of the worms and reproducible lifespan data. In this section, we discuss studies that were conducted to validate our microfluidic approach for lifespan measurement in *C. elegans*. Specifically, (i) we compared the lifespan data for animals cultured and maintained in our microfluidic devices versus those maintained on agar plates, (ii) we compared the stress induced in the liquid culture environment of the microfluidic device to that on agar plates, (iii) we measured lifespan of mutants with known aging pathways, and (iv), we tested efficacy of RNAi interventions in the device.

#### Comparison of *C. elegans* lifespan in device and on agar

To evaluate if our optimized NemaLife device generates *C. elegans* lifespan data consistent with that of standard agar plate assays we conducted parallel lifespan analysis of young adults using established protocols. Figure 5a shows that housing worms in NemaLife does not significantly alter lifespan (p>0.11). The mean (st. dev.) and maximum lifespan from three replicates on agar were 17.5 ± 3.8 and 24 days respectively. Likewise, the mean (st. dev.) and maximum lifespan in the microfluidic device were 17.2 ± 3.4 and 24 days, respectively (see Table S2 for actual data).

**Fig 5.**
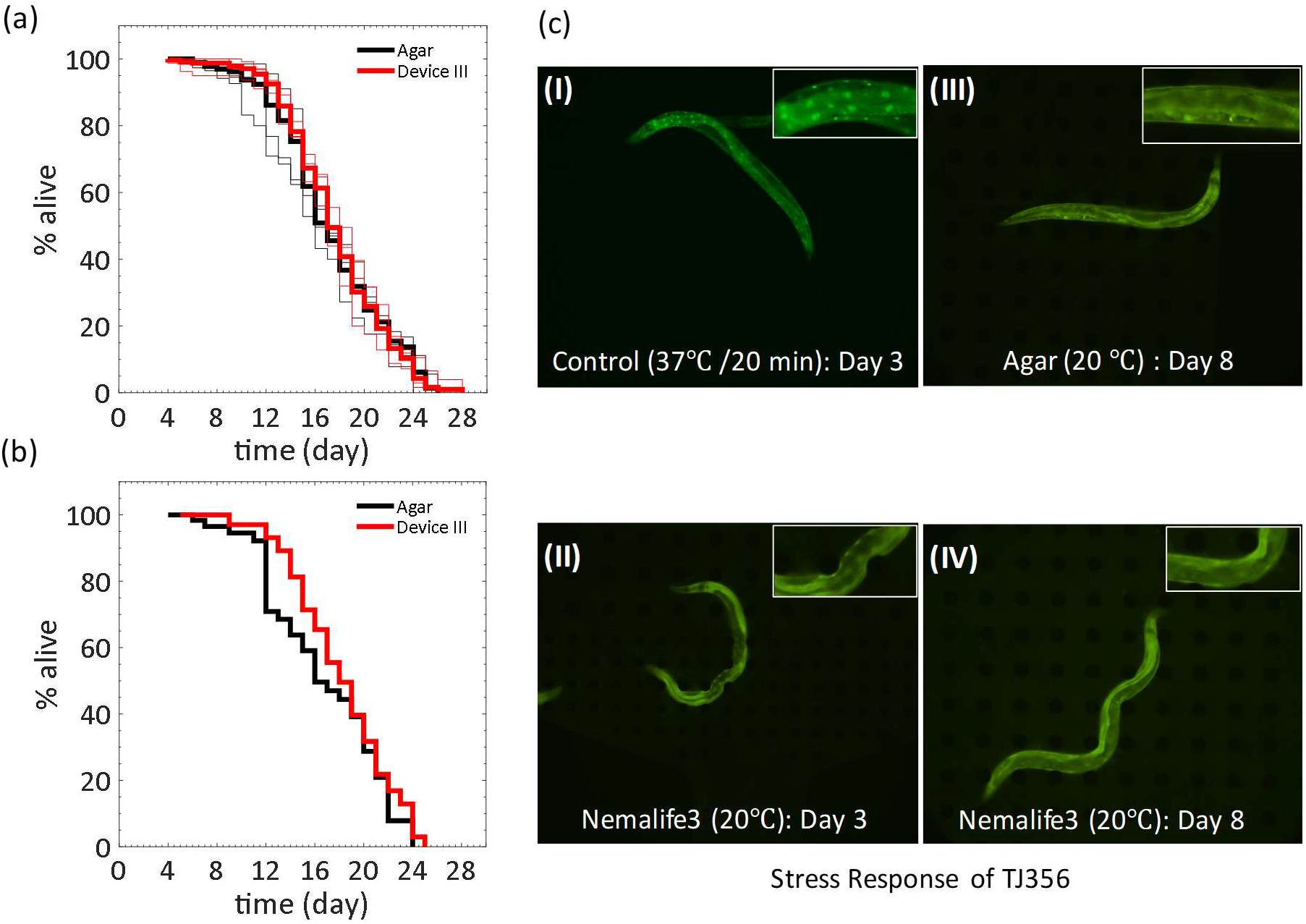
Lifespan measured in the NemaLife device is similar to that animals reared on agar plates: (a) Lifespan of wild type *C. elegans* evaluated on agar plates and in the microfluidic lifespan device. Thin solid lines represent independent trials and the thick solid line represents the combined lifespan of the trials. Lifespan evaluated in microfluidic device is consistent with the agar plate assay (p=0.11 for trial-1; p=0.13 for trial-2 and p=0.22 for trial-3) at 20 °C. Sample size in agar/microfluidic device -trial-1: 109/81; trial-2: 159/135; and trial-3: 70/112. (b) Lifespan of a transgenic strain TJ356 with a Pdaf-16::GFP stress reporter on agar plates and in the microfluidic device (p=0.16). Sample size is 71/112 (agar/microfluidic device) at 20 °C. (c) Fluorescent imaging of DAF-16::GFP nuclear localization of live worm in the microfluidic device. (i) Confirmation of the DAF-16::GFP localization in the nucleus by incubating the worm at 37°C for 20 min and (ii) no sign of accumulation in a day 3 (from hatching) when freshly loaded into the device from agar plate at 20 °C, (iii) and (iv) are images of 8-day old animal (from hatching) on agar plate and in microfluidic device at 20 °C respectively. n = 15. Insets show zoomed-in fluorescence images.

Although, we find that the lifespan curves are in good agreement between NemaLife and agar assays, we find significant differences in loss of animals. On agar with wild-type animals, we found 5 – 30% animal loss. It is common to lose animals on agar plates due to (i) crawling up along the side wall and death from desiccation, (ii) burrowing into the soft agar which precludes scoring, and (iii) bagging (internal hatching). Animal loss due to desiccation and burrowing depends on the type of strain, sex, mutation, and intervention used, which sometimes may account for approximately 50 % of the total animal population^50^. As death events due to desiccation and burrowing cannot be scored, whether agar-based lifespan is a measure of a selected subset of a population becomes a question. Moreover, loss of animals increases the number of worms needed per experiment as loss must be anticipated.

We find that NemaLife eliminates the incidences of animal loss from desiccation and burrowing due to culturing in an enclosed microfluidic chamber. We found 0 − 6 % animal loss in NemaLife which was due to washing mistakes (human error for not plugging the loading port which lacks a sieve channel or when high pressure is applied for fluid flow and animals close to the sieve channel squeeze out). Animal loss/censoring from bagging was 1 − 6% in NemaLife compared to about 2 − 10% on agar plates.

We also evaluated whether worm culture in the liquid environment of the microfluidic device induces environmental stresses such as starvation in the animals. To assess this, we chose a strain that expresses the stress reporter DAF-16::GFP^51^, which exhibits DAF-16 nuclear localization under caloric restriction, heat or oxidative stresses^52–54^. We find that the strain harboring this reporter (TJ356) exhibits similar lifespan on both agar plates and in the microfluidic device, indicating that culture in the liquid environment of the microfluidic device does not induce deleterious effects on lifespan of the reporter strain (Fig. 5b).

We then performed fluorescence imaging to assess DAF-16 localization. As a test of the reporter we exposed the animals in the device to 37 °C for 20 mins and find induction of DAF-16 localization (see the bright spots in the inset image of Fig. 5c-i). In the device at 20 °C, day 3 and day 8 animals, or on agar at day 8 animals did not exhibit such DAF-16::GFP localization (see images in Fig. 5c-ii, iii, iv). Thus, although animals cultured in our microfluidic device can induce stress responses similar to those cultured on agar, the standard growth conditions we use do not elicit daf-16-dependent stress responses.

Over the course of the study, we conducted 20 separate lifespan assays of WT worms in the NemaLife device (Fig. 6), allowing us to account for seasonal changes in laboratory environments. We find that variation in lifespan is limited to 13%, a level of replicate variation comparable to that found in lifespan assays conducted in LSM technolology^50^.

**Fig 6.**
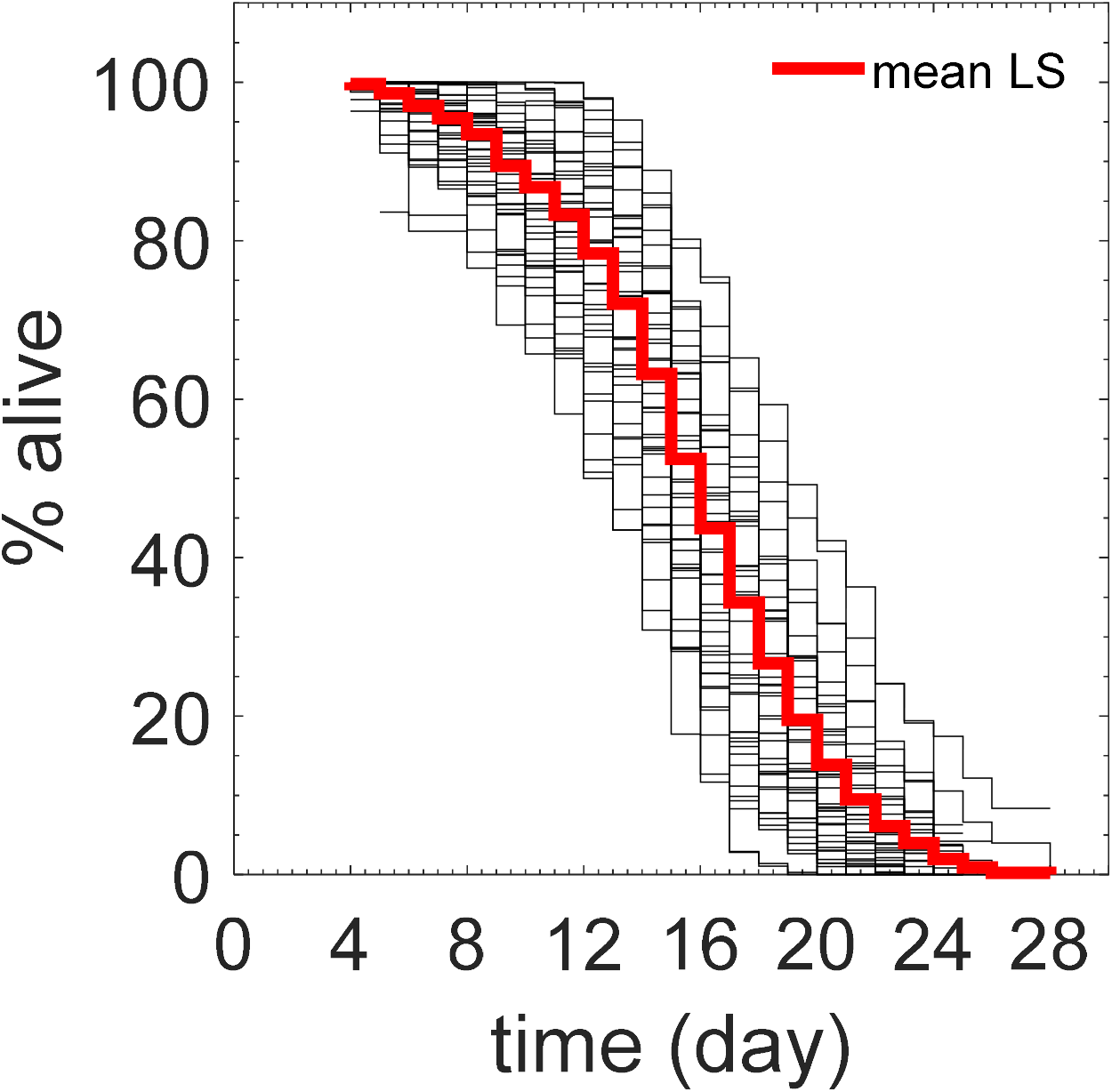
Natural variation in lifespan of wild-type *C. elegans* scored in NemaLife. Lifespan of 20 trials of wild-type animals cultured and scored in NemaLife. Experiments were conducted randomly at different times indicative of seasonal variations in the laboratory environment. Thin black lines are the lifespan curves for individual batch of animals, thick red line represents the mean lifespan derived from the set of 20 experiments. Mean lifespan (95 % C.I.) 13.4 – 15.2 days and maximum lifespan 21.6 – 24.3 days (Range of median lifespan 11 – 19 days and maximum lifespan 17 – 27 days). n = 60 − 150 animals.

#### Lifespan studies using mutants and RNAi interventions

To further test our NemaLife device, we sought to replicate phenotypes of well characterized long-lived and short-lived mutants. As a starting point, we chose established long-lived genetic mutants: insulin signaling mutants *daf-2(e1370)* (insulin receptor reduction of function mutation^55^)*, age-1(hx546)* (phosphatidylinositol-3-OH (PI3) kinase reduction of function mutation^56^) and eating-impaired dietary restriction mutant *eat-2(ad1116)*^57^. We also tested the short-lived, insulin signaling mutant, *daf-16(mgDf50)*, lacking the FOXO transcription factor homolog^58,59^.

Consistent with previous reports, *daf-2, eat-1* and *age-1* mutants exhibited robust extension of maximum lifespan (87%, 33-50%, and 17%, respectively) in the microfluidic environment (Fig. 7a,b; see Table S2 for actual data). Similarly, lifespan analysis of short-lived, *daf-16* mutants is consistent with previous studies (20% reduction in maximum lifespan). We note that we found the maximum lifespan of *daf-2* to be 44 days, which demonstrates that the microfluidic culture environment can adequately support long duration longevity studies in *C. elegans*.

**Fig 7:**
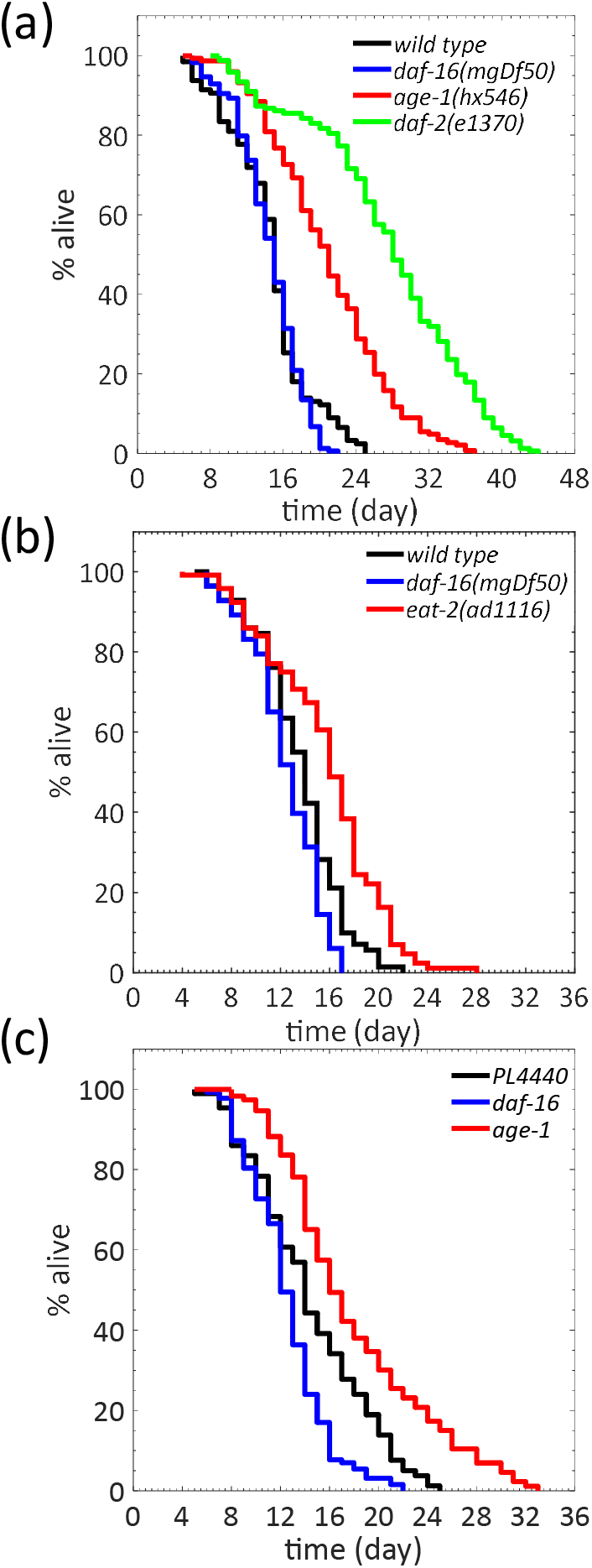
Mutant and RNAi screens in the NemaLife device. (a) Lifespan of long lived mutants *daf-16(mgDf50), daf-2(e1370)* and *age-1(hx546).* 2 trials and n > 160 for *daf-2*, 3 trials and n > 150 for *age-1*, 3 trials and n > 158 for *daf-16* and 3 trials and n > 150 for wild type. (b) Lifespan extension of *eat-2(ad1116)* establishes lifespan device for its suitability to carry out dietary restriction (DR) experiments. 3 trials and n > 112 for *eat-2*, 3 trials and n > 76 for *daf-16* and 3 trials and n > 86 for wild type. (c) RNAi efficacy establishes the ability of the lifespan device to capture the on-chip genetic modification. For RNAi efficacy fragments targeting *daf-16* and *age-1* were inserted into feeding vector L4440. Wild type *C. elegans* of day 3 were used in the experiment. n > 128 for *daf-16*,n >110 for *age-1* and n > 81 for empty vector. N=3 repeat trials. Food: 100 mg *E. coli* OP50/mL S complete, feeding frequency: once/day.

We also tested whether rapid genetic screens through the use of RNAi can be pursued in the NemaLife device. In *C. elegans* RNAi knockdown can be achieved by feeding them bacteria that harbor clones expressing specific double stranded RNAs^60,61^. In Fig. 7c we show the survival curves from RNAi intervention studies including the empty vector control PL4440, which does not have any fragment cloned into it. Compared to this control, we found 25% extension and 8 % reduction in lifespan of *age-1* and *daf-16* respectively (3 replicates), suggesting that the NemaLife microfluidic environment is conducive to RNAi studies. We conclude that NemaLife can both support long term culture of *C. elegans* and document longevity outcomes that parallel those reported on agar plates.

### C. Scoring healthspan measures in *C. elegans*

In addition to lifespan measurement, the transparency and shallow depth of the PDMS worm-habitat chamber offers the opportunity to evaluate physiological measures and fluorescent biomarkers of healthspan in *C. elegans*. In this study, we focused on manual scoring of pharyngeal pumping and stimulus-induced forward and reversal speed. The capacity to score these phenotypic measurements across lifespan through transparent PDMS completes the NemaLife capability as a device for healthy aging investigations in *C. elegans*.

#### Pharyngeal pumping

The pharynx of *C. elegans* is a heart-like organ that uses rhythmic contraction and relaxation to facilitate bacterial uptake^62^. Pharyngeal pumping rates depend on several factors, such as food availability and environmental quality, and significantly decline with age, making it an attractive physiological marker for evaluating *C. elegans* health status^63,64^. Despite the significance of pharyngeal pumping as a physiological marker of aging and sensitivity to environmental conditions, none of the microfluidic lifespan devices reported to date scored this healthspan measure (see Table 1). Instead, specialized devices have been used for measuring pharyngeal pumping^65^.

Using our NemaLife platform, we confirm previous reports of age-dependent reduction of pharyngeal pumping in wild-type *C. elegans* (Fig. 8). Additionally, we observe temporal changes in pharyngeal pumping rates throughout various life stages. In animals maintained within NemaLife, we report that pharyngeal pumping rates increase up to the end of the reproductive period, reaching a maximum of ≈ 283 cycles/min on day 8. Pumping rates decrease gradually starting from day 10, reaching ≈ 117 cycles/min on day 20, at which point, degeneration of the pharynx makes it difficult to score pumping frequency. Maximum pharyngeal pumping rates are similar to that reported on agar^63^, however, the rate of age-dependent decline in the pharyngeal pumping is relatively slow in the microfluidic device compared to agar. Old animals (day 15) grown on agar exhibit a significant reduction in pharyngeal pumping rate (20-30 cycles/min^63,64^ while animals housed in the microfluidic device maintain a relatively high pharyngeal pumping rate (150-160 cycles/min) in late life. The reason for longer maintenance of pumping rate in the microfluidic device is not clear, however, this feature may be advantageous for enhanced uptake of compounds in pharmacological assays.

**Fig 8:**
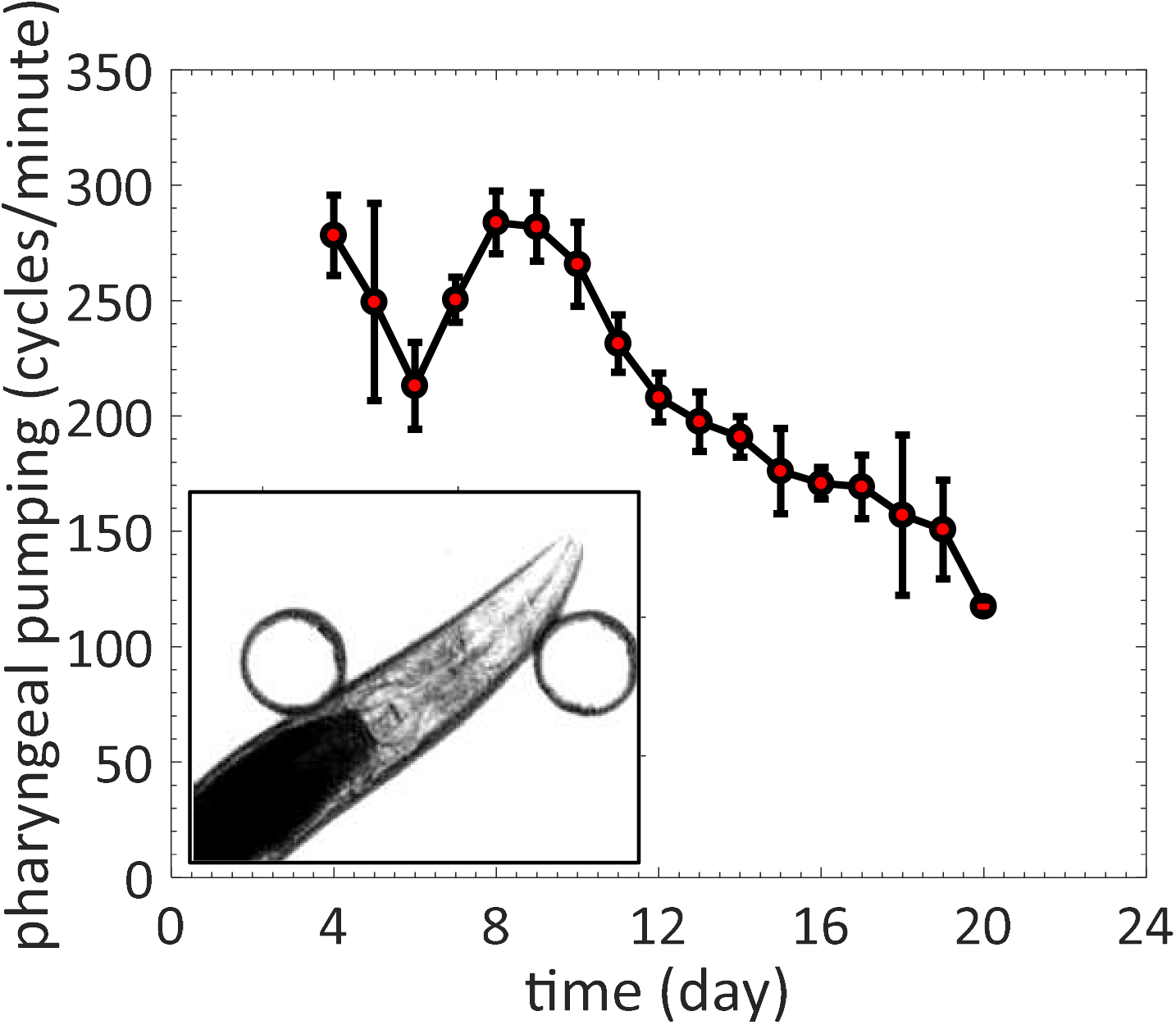
Age-associated decline in pharyngeal pumping rate as measured in the NemaLife device. (a) Pharyngeal pumping in cycles per minute for wild type *C. elegans* over the course of its lifespan (n=10 at each time point). Error bar is the standard deviation. Food: 100 mg *E. coli* OP50/mL S complete, feeding frequency: once/day. Inset shows the pharynx of an animal inside a chamber.

#### Stimulated locomotion

Locomotory vigor is commonly used as a healthspan measure in *C. elegans*^63,64,66^. Forward crawling of *C. elegans* is accompanied by pauses and reversals at a speed and frequency that is dependent upon food availability and the crawling environment. As a result, temporal fluctuations in locomotion make it difficult to evaluate true crawling speed. Extended tracking and long-term analysis of time-lapse images is required to properly assess forward locomotion dynamics. Reversals are observed during natural locomotion, but are also important in escape response to gentle touch in which the worm quickly reverses and suppresses head movement^67–70^. Spontaneous reversals are usually short episodes between consecutive forward crawling bouts. Reversal behavior is thus a strong indicator of neuro-muscular function^69^, that can be scored reliably in a short observation time.

On agar plates, reversal is induced by applying gentle touch to the animal with eyelash or by prodding the worm with worm pick or by simply tapping the plate^71,72^. Here, we replicate the gentle touch stimulus in the microfluidic device with a hex key to induce stimulated reversals (see **SI movie 4**). We induced reversals mechanically by gentle tapping on the top surface of the device 3 times at a location slightly away from the worm pharynx. Stimulus is transferred to the worm as mechanical vibration through the pillar and the fluid. This stimulus-induced locomotory response cannot be evaluated in prior microfluidic devices that use swimming worms and is unique to the NemaLife device.

Figure 9a is a time series of instantaneous speed of a day 4 (from hatching) wild-type animal calculated from the behavioral episode during the gentle-touch stimulus in the NemaLife device. Stimulated responses generally involve an initial reversal (first reversal), a change in direction, followed by a final forward movement. In nearly all cases, we observe animals immediately respond by exhibiting a first reversal followed by a turn. Occasionally, we observe brief interruptions in forward crawling motion that appears to be independent of the stimulus. Inset of Fig. 9a shows the average (n = 5 animals) speed calculated from the different modes of crawling. As expected, the first reversal shows the highest speed.

**Figure 9.**
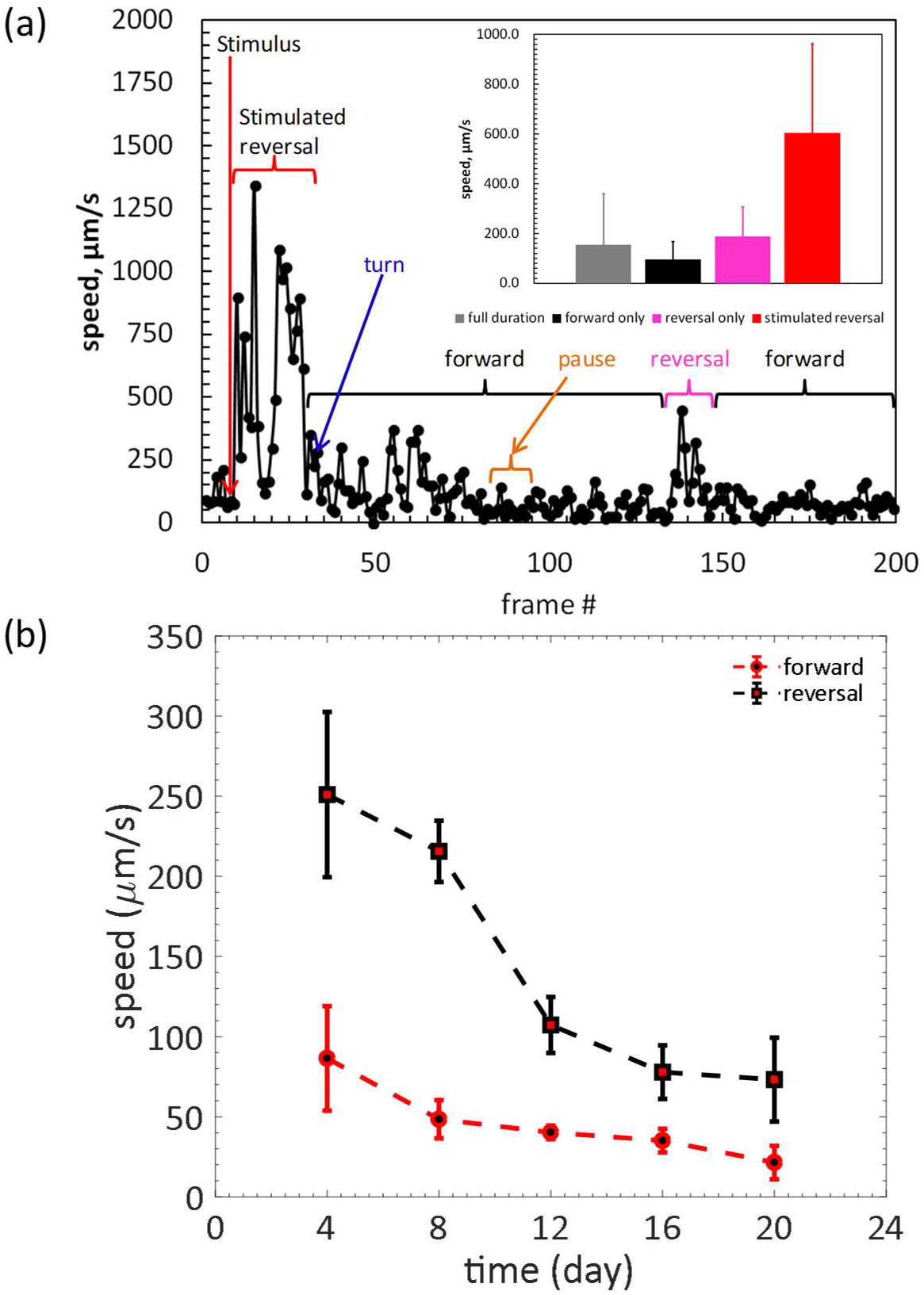
Stimulated reversal speed and age-associated locomotory decline in wild-type *C. elegans* as measured in the NemaLife device. (a) A representative locomotion episode of a day-5 *C. elegans* over 40 seconds duration. Each filled circle represents instantaneous speed between two consecutive frames determined by manually tracking the vulva of the animal. Tap stimulus was applied (frame # 10, red arrow) to induce reversal. Animal executed reversal with a very high speed followed by second reversal. Animal turns (blue arrow), crawls in forward direction (black brackets) for relatively long time with pause (amber bracket). Naturally occurring reversal period is shown by the pink bracket. Average speed of the full episode, only forward (including pauses), only reversal and stimulated reversal are shown in the inset. Error bars are standard deviations. n = 5. (b) Decline in forward and first reversal speed as wild type *C. elegans* ages in the micropillar arena. Error bar is the standard deviation. n > 10, food: 100 mg *E. coli* OP50/mL S complete, feeding frequency: once/day.

Using our NemaLife platform, we show that both reversal and forward speed varies significantly with age and we find that reversal speed is always greater than forward speed in worms of all ages (Fig. 9b). Most importantly, the rate of decline in reversal speed is accelerated at the end of reproduction, specifically 50%, compared to a substantially smaller decline in stimulated forward speed (17%). Interestingly, the decline in the stimulated reversal speed is correlated with an equally rapid decline in survival rates following the reproductive period. These observations underscore the importance of expanding lifespan assays to include the evaluation of additional health span metrics. Overall, our NemaLife device enables measurement of stimulated reversal speed as a novel biomarker for aging and healthspan, which couples with other health measures to provide a powerful platform for analysis of *C. elegan*s healthspan and lifespan.

## IV. Conclusions

We successfully demonstrated that *C. elegans* can be effectively maintained in our NemaLife microfluidic device across its lifespan without using chemicals (progeny-blocking drugs, antibacterial agents, antifungal compounds, etc.) in an environment that recapitulates longevity on agar plates. Micropillars in the microfluidic device enable the animals to maintain natural crawling gaits and eliminate swim-induced fatigue. The longevity outcomes of “cornerstone” mutants with altered insulin-like signaling or dietary restriction pathways grown on agar are reproduced in the device; RNAi can also be executed. The capacity for manual injections with syringes and investigator scoring provides control over spurious data and unintended events. Altogether, our NemaLife device is a simple and low-cost means to obtain reliable lifespan and physiological data. It can be developed into a fully automated platform, including progeny removal, feeding, drug delivery, and scoring phenotypes using automated pump systems and the appropriate software. Efficient progeny removal without clogging is a significant advantage if the NemaLife is integrated into an automated platform. Thus the device has a significant potential both as a low cost device to be used in many labs and as a key component of a fully automated platform.

Easy fluid exchange ability in a confined space provides unprecedented temporal control over the animal habitat. This temporal control allows addition or withdrawal of multiple stimuli, enabling researchers to design sophisticated multi-step experiments without increasing technical difficulty or adding significant time devoted to such assays. In addition, transparent PDMS allows both brightfield and fluorescent imaging of live worms, facilitating life-long observation of cellular and sub-cellular components. Therefore, the health of aging worms can be measured using multiple parameters (pumping rate, velocity), virtually from the womb to tomb. A potential benefit of the device is that progeny and effluent (containing any chemicals such as, pheromones etc.) might be collected for downstream analysis. With automation of fluidics and scoring, we anticipate that the simplicity of our method, combined with unprecedented capacity to temporally manipulate the environment of the animals and record multidimensional healthspan measures, will enable large-scale parallelized cross-sectional and longitudinal aging experiments.

## V. Experimental Procedures

### Worm culture

All animals were cultured on 60 mm petri dishes containing nematode growth medium (NGM) at 20°C before loading into the microfluidic chamber. The NGM filled petri dishes were seeded with 300-400 μL of bacteria *Escherichia coli* OP50 and incubated for 48 hours at 20°C. For age synchronization, 20-25 gravid adults were placed on seeded plates to lay eggs for 2-4 hours. After eggs were laid, the animals were removed from the plates, eggs were incubated for 60-72 hours. The day the eggs were laid was scored as day 0. In this study, we used wild-type Bristol (N2), GR1307[*daf-16 (mgDf50)I*]*, CB1370[daf-2 (e1370)III*], TJ1052[*age-1(hx546)II*], DA1116[*eat-2(ad1116)II*] and TJ356[*zIs356* [*daf-16*p::*daf-16a/b*::GFP + *rol-6(su1006)]IV*]. Wild-type (N2), *daf-16*, and *daf-2* mutants were received from the *Caenorhabditis* Genetics Center (CGC); *age-1* and *eat-2* strains were kindly provided by the Driscoll lab.

### Device fabrication and preparation

All microfluidic devices were fabricated in poly(dimethyl)siloxane (PDMS) using soft lithography^73^. A mold was fabricated using two-step SU-8 photolithography such that the chamber height is ≈ 100μm and the micropillar height is ≈ 75 μm, as described previously^49^. A 4-6 mm thick PDMS (Sylgard 184 A and B, 1:10 by weight, Dow Corning) layer was casted on to the mold and the inlet/outlet holes were punched with a 1 mm hole puncher. The PDMS device was then bonded on a glass surface irreversibly and rendered hydrophilic by plasma treatment (Harrick Plasma inc.). Before using the device for lifespan experiments, the device interiors were filled with 70% ethanol for 5 minutes to sterilize them. Subsequently, the device was rinsed 4-5 times with S-complete solution. Devices were then treated with 5 wt% Pluronic F127 (Sigma-Aldrich) for 30 minutes to prevent protein and bacterial build-up^33^. In addition, Pluronic treatment also assists with removal of air bubbles if any are trapped. After incubation, excess Pluronic was removed by washing with S-complete. The Pluronic-treated devices were stored in moist petri dishes at 20°C for immediate use and at 4°C for future use.

### Food preparation

*E. coli* OP50 was used as the bacterial food source for worms grown on both NGM and maintained within the devices. Bacterial suspension of 100 mg mL^−1^ in S-complete solution corresponding to ≈ 10^9^ bacteria/mL was used for lifespan assays unless otherwise noted. E. *coli OP50* was grown overnight at 37°C in standard LB broth. Bacterial suspensions of 100 mg mL^−1^ were prepared by centrifuging 500 mL of overnight bacterial culture and resuspending the pellet in S-complete. Concentrated OP50 was stored at 4°C for subsequent use, up to two weeks.

### Bacteria preparation for RNAi studies

Engineered bacteria expressing double-stranded RNA (dsRNA) were obtained from the Driscoll Lab and used for testing the RNAi efficacy in the device. Fragments designed for targeting *daf-16* and *age-1* were cloned into the L4440 feeding vector and the resulting plasmids were transformed into the HT115 (DE3) using standard protocols^61^. Bacteria with empty L4440 vector (no cloned fragments) were used as the negative control for all experiments involving RNAi. Bacterial colonies were grown on LB agar supplemented with 50 µg/ml carbenicillin for 48 hours at 37°C. Fresh plates were made each week.

Single colonies of bacteria were picked and grown in culture flasks with shaking at 200 rpm for 16 hours in sterile LB broth with 50 µg/ml ampicillin at 37°C. For induction, 0.4 mM Isopropyl β-D-1-thiogalactopyranoside (IPTG) was added for 2 hours at 37°C while shaking. At 18 hours, additional IPTG was added for a final concentration of 1 mM. Concentrated bacterial solutions were prepared each day using procedures described above. Final IPTG concentration was maintained at 1 mM in the food solution.

### Fluorescence imaging

We imaged the *C. elegans* strain containing stress reporter gene *zls356IV* inside the device without immobilizing them using a Nikon Ti microscope at 10X magnification. Movies were captured using fast time lapse imaging with a camera (Zyla 5.5 sCMOS from Andor Inc.). For imaging worms cultured on agar plates, worms were loaded into a fresh chamber and imaged immediately. Movies were analyzed using ImageJ (NIH)software.

### Scoring animal death

Worms were counted manually and lifespan was scored daily. An animal was scored as dead if it failed to respond to (i) gentle flow of fluid throughout the chamber or (ii) gentle tapping of the device by a 3/8” Allen key. If there is no movement in the pharynx or in the tail 1 minute after the stimulus has been applied, we scored the animal as dead. Each death event was scored as 1 and unaccounted deaths (missing, washing error, matricides) were scored as 0. A lifespan curve (Kaplan-Meier) was then generated. Kaplan-Meier curves and Log-Rank statistics were generated using the Statistics Toolbox in MATLAB.

### Locomotory measures

Reversal and forward speed were scored after a gentle tap on the PDMS device with a 3/8” Allen key, at a location close to the tip of the pharynx. Only the initial, instantaneous reversal following stimulation were scored. Continuous frames of a spontaneous start of reversal and end of reversal were taken as a complete reversal episode. Reversal episodes with a pause, stop, or intermittent reversal were not included. Continuous forward locomotion until a spontaneous reversal was scored to determine forward speed. Images were captured using SVSi streamview camera at 10 frames/second. Locomotion speed was calculated by tracking the displacement of the pharynx in time. The maximum speed in an episode (forward/reversal) of a worm was recorded as the maximum reversal/forward speed.

### Pharyngeal pumping

Movies of 10 individual worms were captured at a rate of 20 frames/second with SVSi streamview camera on a Zeiss stereo microscope at 5X magnification, 15 minutes after the addition of food to the device. Complete cycle time of contraction/retraction of the isthmus/terminal bulb of pharynx was manually computed using ImageJ (NIH). The number of pharyngeal pumping cycles were counted over a 10 second period for each single animal and then reported as cycles per minute.

### Data analysis

All the survival analyses were conducted in MATLAB (Mathworks, R2014b). Log-rank (Mantel-Cox) test was used to compare survival between treatment groups. Two-sample t-test was used to compare growth of the worm and crawling kinematics in the optimization study^74^.

## Acknowledgements

This work was partially supported by funding from NIH (Grant Nos. R21 AG050503, R01 AG051995-01A1), NASA (Grant No. NNX15AL16G), NSF CAREER (Grant No.1150836) and BBSRC (Grant No. N015894). We are grateful to Jen Hewitt and Swastika Bithi for useful discussions. We thank Caenorhabditis Genetics Center (CGC) for providing strains, which is funded by NIH Office of Research Infrastructure Programs (P40 OD010440). We also thank members of the Driscoll lab for useful discussions.

## Electronic Supplementary Information (ESI)

Supplementary Materials is uploaded as a separate file. List of the items are

Supplementary Figure.1. Changes in body size of *C. elegans* cultured in the NemaLife device.

Supplementary Table 1. Measured dimensions of the micropillar geometries used in the study for device optimization.

Supplementary Table 2. Summary of survival analysis of the strains and culture methods used in the study.

**Supplementary Video 1: Overall culture (adult *C. elegans* population and their progeny) in a chamber before washing.** A NemaLife chamber with 33 animals, their progeny (grown over 24 hours) and eggs. This video explains an exaggerated scenario before washing. The chamber was purposefully loaded with approximately two times larger population than the chamber capacity, so that many progenies is available to wash and visualize. The movie plays at a speed of 10 fps.

**Supplementary Video 2: Illustration of flow induced progeny separation.** Flow is induced from the loading port with a hand operated syringe. Flow was paused approximately after each 200 µL. The movie plays at a speed of 10 fps.

**Supplementary Video 3: Addition of food (100 mg/mL *E. Coli* OP50).** Food is added through the loading port using a hand operated 1 mL syringe. The movie plays at a speed of 10 fps.

**Supplementary Video 4: Stimulated reversal.** Reversal is induced reproducibly by tapping the device gently with a hex key. Tap is applied between frame 24 and 25 in this movie which is seen as dark shadow at the bottom left corner of the frame. The movie plays at a speed of 10 fps.

## Supplementary information

**Figure S1:**
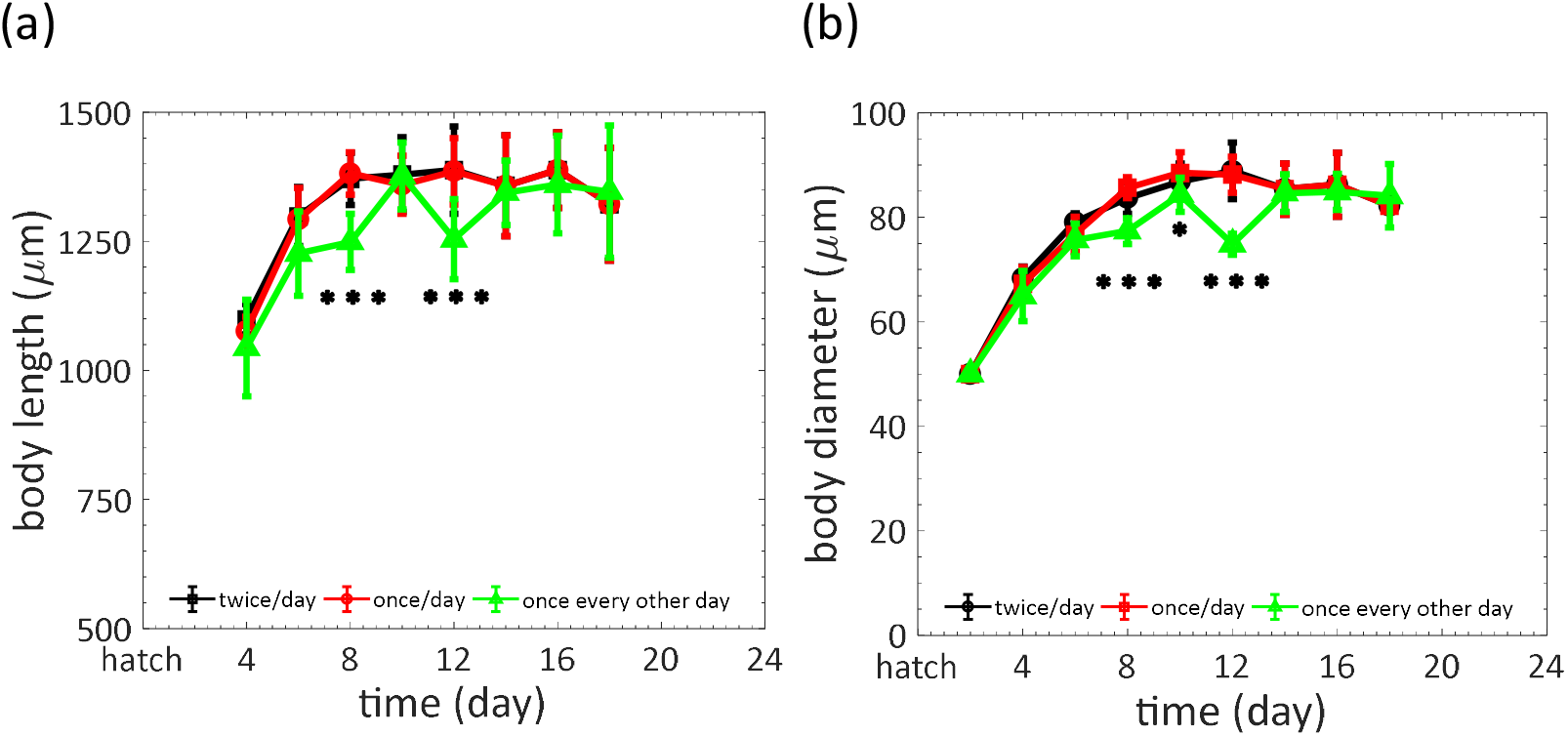
Changes in body size of *C. elegans* cultured in the NemaLife device. (a) Average worm length of wild type *C. elegans* corresponding to the lifespan of main text figure 2(a). Error bars are standard deviations. n>10.(b) Average worm diameter of wild type *C. elegans* corresponding to the lifespan of main text figure 2a. Error bars are standard deviations. n>10*P-value (once/day and once every other day) <<0.01, *** P-value (once/day and once every other day) <<0.0001.

**Table S1:** Measured dimensions of the micropillar geometries used in the study for device optimization.

| **Device** | **Pillar diameter,**<br>**a ($\mu\text{m}$)** | **Pillar height,**<br>**h ($\mu\text{m}$)** | **gap between**<br>**pillars, s ($\mu\text{m}$)** | **Clearance,**<br>**c ($\mu\text{m}$)** |
| --- | --- | --- | --- | --- |
| Device-I | $38.0 \pm 0.4$ | $79.7 \pm 2.9$ | $62.4 \pm 2.9$ | $23.1 \pm 2.9$ |
| Device-II | $52.0 \pm 2.1$ | $77.1 \pm 2.0$ | $78.0 \pm 2.2$ | $27.0 \pm 0.8$ |
| Device-III | $61.3 \pm 1.6$ | $78.4 \pm 1.3$ | $98.5 \pm 1.7$ | $28.4 \pm 3.1$ |

**Table S2:** Summary of survival analysis of the strains and culture methods used in the study.

|  | Type of food | Culture media | Genotype | Sample size, N | Average lifespan (day) |  | median lifespan (day) | Max lifespan (day) |
| --- | --- | --- | --- | --- | --- | --- | --- | --- |
|  |  |  |  |  | mean | s.d. |  |  |
| Agar Vs NemaLife | *E. Coli* OP50 | Agar plate | wild type (N2-bristol) | 109 | 15.95 | 5.01 | 15 | 25 |
|  | *E. Coli* OP50 | NemaLife | wild type (N2-bristol) | 81 | 17.75 | 5.18 | 16 | 27 |
|  | *E. Coli* OP50 | Agar plate | wild type (N2-bristol) | 159 | 17.48 | 3.80 | 16 | 24 |
|  | *E. Coli* OP50 | NemaLife | wild type (N2-bristol) | 135 | 17.16 | 3.40 | 16 | 24 |
|  | *E. Coli* OP50 | Agar plate | wild type (N2-bristol) | 70 | 17.29 | 4.67 | 16 | 24 |
|  | *E. Coli* OP50 | NemaLife | wild type (N2-bristol) | 112 | 16.70 | 3.46 | 16 | 23 |
| Stress evaluation | *E. Coli* OP50 | Agar plate | TJ 356[daf-16(zIs356 IV)] | 71 | 16.44 | 4.8 | 15 | 23 |
|  | *E. Coli* OP50 | NemaLife | TJ 356[daf-16(zIs356 IV)] | 112 | 18.22 | 3.92 | 18 | 24 |
| RNAi efficacy | *E. Coli* OP50 | NemaLife | wild type (N2-bristol) | 127 | 14.3 | 4.4 | 13 | 25 |
|  | Empty vector PL4440 | NemaLife | wild type (N2-bristol) | 88 | 14.4 | 4.8 | 13 | 24 |
|  | daf-16 fragment cloned into HT115 | NemaLife | wild type (N2-bristol) | 135 | 12.6 | 3.2 | 11 | 21 |
|  | age-1 fragment cloned into HT115 | NemaLife | wild type (N2-bristol) | 116 | 17.5 | 5.9 | 15 | 32 |
|  | empty vector PL4440) | NemaLife | wild type (N2-bristol) | 131 | 11.9 | 3.4 | 11 | 21 |
|  | daf-16 fragment cloned into HT115 | NemaLife | wild type (N2-bristol) | 137 | 11.7 | 3.4 | 11 | 19 |
|  | age-1 fragment cloned into HT115 | NemaLife | wild type (N2-bristol) | 150 | 14.3 | 5.2 | 13 | 31 |
|  | empty vector PL4440 | NemaLife | wild type (N2-bristol) | 102 | 12.8 | 3.4 | 12 | 18 |
|  | daf-16 fragment cloned into HT115 | NemaLife | wild type (N2-bristol) | 123 | 11.5 | 3.3 | 11 | 17 |
|  | age-1 fragment cloned into HT115 | NemaLife | wild type (N2-bristol) | 118 | 13.7 | 3.9 | 13 | 24 |
| Lifespan influencing mutants | *E. Coli* OP50 | NemaLife | wild type (N2-bristol) | 157 | 14.8 | 4.0 | 15 | 27 |
|  | *E. Coli* OP50 | NemaLife | *age-1 (hx546)* | 174 | 19.6 | 4.5 | 18 | 33 |
|  | *E. Coli* OP50 | NemaLife | *daf-2 (e1307)* | 163 | 18.0 | 4.4 | 17 | 31 |
|  | *E. Coli* OP50 | NemaLife | wild type (N2-bristol) | 86 | 13.5 | 3.5 | 13 | 21 |
|  | *E. Coli* OP50 | NemaLife | *daf-16 (mgDf50)I* | 84 | 12.5 | 2.9 | 12 | 16 |
|  | *E. Coli* OP50 | NemaLife | *age-1 (hx546)* | 123 | 15.3 | 4.8 | 15 | 27 |
|  | *E. Coli* OP50 | NemaLife | *age-1 (hx546)* | 152 | 20.8 | 6.5 | 23 | 36 |
|  | *E. Coli* OP50 | NemaLife | wild type (N2-bristol) | 128 | 14.6 | 4.5 | 14 | 24 |
|  | *E. Coli* OP50 | NemaLife | *daf-16 (mgDf50)I* | 168 | 14.5 | 3.5 | 14 | 21 |
|  | *E. Coli* OP50 | NemaLife | *daf-2 (e1307)* | 160 | 27.8 | 8.7 | 28 | 44 |
|  | *E. Coli* OP50 | NemaLife | wild type (N2-bristol) | 86 | 13.5 | 3.5 | 13 | 21 |
|  | *E. Coli* OP50 | NemaLife | *daf-16 (mgDf50)I* | 84 | 12.5 | 2.9 | 12 | 16 |
|  | *E. Coli* OP50 | NemaLife | *eat-2 (ad1116)II* | 123 | 15.3 | 4.8 | 15 | 27 |

